# YAP/TAZ inhibition refines TGF-β signaling to prevent laryngeal fibrosis

**DOI:** 10.64898/2026.05.18.726086

**Authors:** Ryosuke Nakamura, Renjie Bing, Hannah Weber, Masayoshi Yoshimatsu, Gary Gartling, Michael J. Garabedian, Ryan C. Branski

## Abstract

Voice disorders affect nearly 20 million Americans and cost more than $13 billion annually. Vocal fold (VF) fibrosis, a major cause of chronic dysphonia, disrupts normal vocal fold vibration by replacing the flexible extracellular matrix with stiff fibrotic tissue. Although TGF-β drives fibrosis, it also activates intrinsic negative feedback mechanisms, including SMAD7 induction and SMAD3 downregulation, to restrain excessive signaling. Broad inhibition of TGF-β or canonical SMAD signaling may disrupt these protective feedback loops and impair normal tissue homeostasis. An ideal anti-fibrotic strategy should differentially target the pro-fibrotic output of TGF-β. Here, we show YAP/TAZ inhibition selectively suppresses pro-fibrotic TGF-β signaling in VF fibroblasts. Pharmacologic inhibition of YAP/TAZ blocked TGF-β-induced fibroblast activation and fibrotic gene expression, while only modestly affecting canonical SMAD feedback responses. Integrated RNA-seq and ChIP-seq analyses demonstrated YAP/TAZ primarily regulate non-canonical TGF-β signaling and pro-fibrotic transcriptional programs. In a rat model of VF fibrosis, YAP/TAZ inhibition reduced nuclear YAP/TAZ localization and attenuated scar formation. Together, these findings identify YAP/TAZ inhibition as a promising therapeutic strategy for VF fibrosis and other fibrotic diseases.

## INTRODUCTION

The vocal folds (VFs) sit at the intersection of the upper and lower airway and play key roles in swallowing, airway protection, and in many species, communication. The VFs withstand extensive environmental and mechanical insults, leading to aberrant tissue structure and function. Voice disorders affect nearly 20 million Americans at a direct cost estimated at over $13 billion annually.^1^ The mucosa of the VFs has a unique extracellular matrix (ECM) enriched with hyaluronic acid. Fibrosis of the VF mucosa is largely intractable and results in replacement of the unique pliable ECM with stiff fibrous proteins, and altered tissue architecture leading to substantive communication impairment. Impaired voice quality is associated with profound social and occupational limitations, and in turn, reduced quality of life and economic productivity.^2,3^ Current therapeutic options, such as surgical intervention, remain limited.

Steroids, broadly employed for benign VF lesions including fibrosis, are also associated with inconsistent outcomes. Identification of relevant therapeutic targets based on molecular pathophysiology is key to developing innovative and effective treatments for VF fibrosis and may apply more broadly to other fibrotic disease processes.^4–6^

Fibroblast activation is a key pathological event leading to fibrosis. Transforming growth factor-β (TGF-β) signaling triggers fibroblast activation, inducing a contractile phenotype and hyper-accumulation of ECM primarily via the SMAD2/3-mediated canonical pathway.^7^ However, beyond its roles in the fibrotic response, TGF-β signaling broadly regulates fundamental cell activities, such as cell proliferation and survival, through both canonical and non-canonical pathways.^8^ Moreover, SMAD2/3 drives intrinsic negative feedback mechanisms, such as SMAD7 upregulation.^9^ Unraveling the complexity of TGF-β signaling in VF fibroblasts holds significant potential for identifying novel therapeutic targets to suppress the fibrotic response while sparing homeostatic TGF-β-mediated processes.

Yes-associated protein (YAP) and transcriptional co-activator with PDZ-binding motif (TAZ) are transcriptional co-regulators that regulate gene expression by working with TEA domain transcription factors (TEADs) to control multiple signaling pathways.^10,11^ Aberrant activation of YAP/TAZ is associated with various pathologies, including fibrosis.^12^ Genetically induced YAP/TAZ activation exacerbated hepatic and renal fibrosis, and recent studies from our laboratory demonstrated TGF-β activates YAP/TAZ in VF fibroblasts in parallel with SMAD2/3 activation.^13–15^ YAP/TAZ inhibition has shown promise in animal models of hepatic, pulmonary, and renal fibrosis.^16^ In malignant mesothelioma cells, a YAP-TEAD4-SMAD2/3 tertiary complex likely contributes to transmission of TGF-β signaling.^17^ However, despite the important roles of YAP/TAZ in fibrosis, the molecular mechanisms underlying their interactions with SMAD2/3 remain unclear.

In this study, we investigated how SMAD2/3 and YAP/TAZ regulate gene expression in VF fibroblasts. Here, we report YAP/TAZ inhibition greatly reduced expression of key fibrotic genes such as *ACTA2* and *FN1*, while having only a moderate impact on negative feedback mechanisms within the TGF-β/SMAD signaling cascade. These findings suggest YAP/TAZ preferentially mediates the pro-fibrotic response among the complex events induced by TGF-β signaling. To further investigate the roles of YAP/TAZ, as well as the therapeutic effect of YAP/TAZ inhibition, we used a multi-omics approach to compare YAP/TAZ and SMAD2/3-dependent gene expression, and tested pharmacological YAP/TAZ inhibition in a rat model of VF fibrosis. These data provide a foundation for improved understanding of the pathophysiology of VF fibrosis and development of novel therapeutic strategies for laryngeal disease associated with fibrosis.

## RESULTS

### YAP/TAZ inhibition reduced expression of key fibrotic genes induced by TGF-β

A human VF fibroblast cell line, HVOX,^18^ was treated with TGF-β1 (10ng/mL) together with increasing concentrations of K-975 to assess the impact of pharmacological YAP/TAZ inhibition on TGF-β-induced fibroblast activation. K-975, an irreversible inhibitor of YAP/TAZ binding to TEADs,^19^ reduced expression of genes involved in tissue contraction (*ACTA2*) and ECM deposition (*FN1*, *COL1A1*, and *LOX*) in a concentration-dependent manner, with statistically significant inhibition observed from 0.3μM to 1μM **(Fig. 1a–d)**. Of note, treatment with 10μM K-975 almost completely blocked TGF-β-induced upregulation of *ACTA2*, *FN1*, and *LOX* **(Fig. 1a, c, d)**. Protein levels of α-smooth muscle actin (αSMA), type I collagen, and fibronectin (encoded by *ACTA2*, *COL1A1*, and *FN1*, respectively) were also decreased by K-975 treatment **(Fig. 1e)**. These data suggest pharmacological inhibition of YAP/TAZ efficiently suppressed gene expression associated with fibroblast activation, even though K-975 does not directly target canonical TGF-β/SMAD signaling.

**Figure 1.**
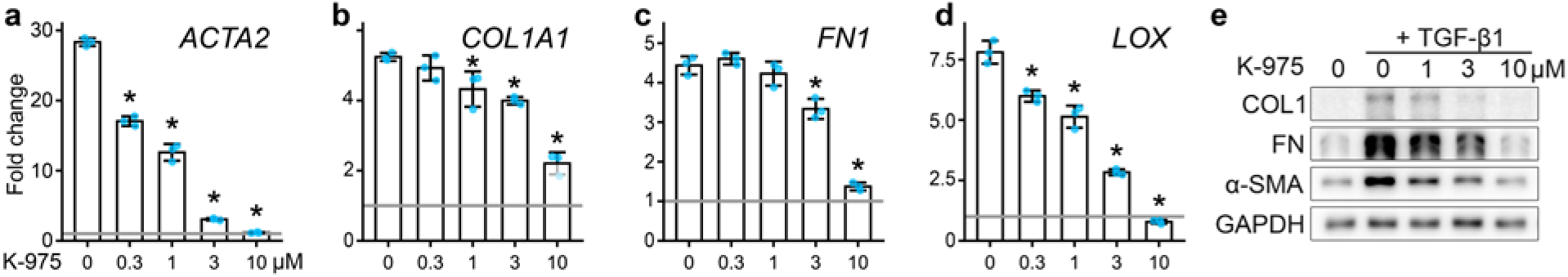
YAP/TAZ inhibition had an anti-fibrotic effect on human VF fibroblasts *in vitro*. A human VF fibroblast cell line, HVOX, was treated with 10ng/mL TGF-β1 +/- 0.3‒10μM K-975 for 24 hours. Expression of genes associated with fibroblast activation was assessed by RT-qPCR (a–d). Fold changes relative to control cells, without TGF-β1 or K-975 treatment, are shown as bar graphs (mean ± SD). Significant difference is indicated with asterisks (*n* = 3, *p* < 0.05, Dunnet test). Western blots show protein expression of alpha-smooth muscle actin (α-SMA), type I collagen (COL1), and fibronectin (FN) (e; whole images are available in Supplementary Material Fig. S1). Glyceraldehyde 3-phosphate dehydrogenase (GAPDH) expression was assessed as an internal control. Representative data from triplicate experiments are shown.

### YAP/TAZ inhibition moderately impacted conventional SMAD2/3 target genes

To assess how YAP/TAZ inhibition effects YAP/TAZ and SMAD2/3 signaling, expression of YAP/TAZ target genes (*CCN1*, *CCN2*, *EDN1*, *THBS1*, and *TGM2*), SMAD2/3 target genes (*SERPINE1* and *SMAD7*),^20–24^ and SMAD2 and SMAD3 were assessed by quantitative real-time polymerase chain reaction (RT-qPCR) in response to TGF-β1 -/+ K-975 treatment. TGF-β1 induced *CCN1*, *CCN2*, *EDN1*, and *THBS1* expression and treatment with 10μM K-975 decreased expression to basal levels **(Fig. 2a–d)**. Since *CCN2* encodes connective tissue growth factor, a major driver of the fibrotic response, and *THBS1* encodes thrombospondin-1, which activates latent TGF-β1, downregulation of these genes likely underlies the anti-fibrotic effect of YAP/TAZ inhibition.^25,26^ *TGM2*, which encodes transglutaminase 2 involved in ECM cross-linking,^27^ was not induced by TGF-β1, but was significantly downregulated by K-975 **(Fig. 2e)**.

**Figure 2.**
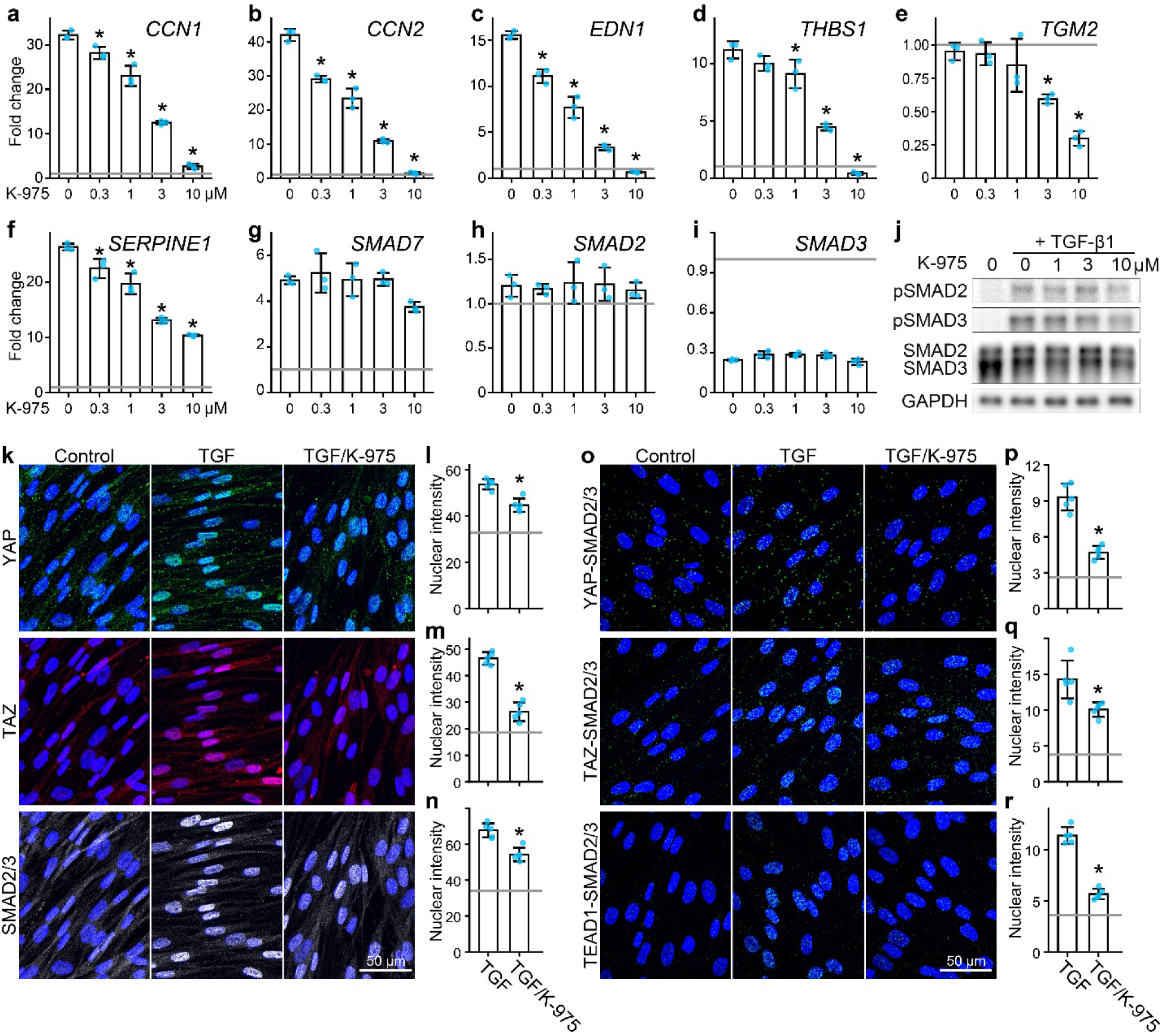
YAP/TAZ inhibition reduced both YAP/TAZ and SMAD2/3 activity in human VF fibroblasts *in vitro*, with modest impact on SMAD2/3 target genes. Human VF fibroblasts were treated with 10ng/mL TGF-β1 +/- 0.3‒10μM K-975 for 24 hours. Expression of target genes for YAP/TAZ (a-e) and SMAD2/3 signaling (f-i) was assessed by RT-qPCR. Fold changes relative to control cells, not treated with TGF-β1 or K-975, are shown as bar graphs (mean ± SD). Significant difference compared to cells treated with TGF-β1 alone is indicated as asterisks (*n* = 3, *p* < 0.05, Dunnet test). Western blots show protein levels of Ser465/467-phosphorylated SMAD2 (pSMAD2), Ser423/425-phosphorylated SMAD3 (pSMAD3), and total SMAD2 and SMAD3 at 12 h after treatment (j; whole images are available in Supplementary Material Fig. S2). Glyceraldehyde 3-phosphate dehydrogenase (GAPDH) expression was assessed as an internal control. Representative data from triplicate experiments are shown. Immunofluorescene images show YAP (green), TAZ (red), and SMAD2/3 (grey) in HVOX cells exposed to 10ng/mL TGF-β1 +/- 10μM K-975 for 12 h (k). Nuclei were counterstained with DAPI (blue). Immunofluorescence signal intensities in the nucleus are shown as bar graphs (mean ± SD; l–n). Significant difference is indicated with asterisks (*n* = 3, *p* < 0.05, Student *t*-test). Proximity ligation assay (PLA) visualized close localization of YAP, TAZ, TEAD1, and SMAD2/3 (o; green: PLA signal; blue: DAPI). Intensities of PLA signals in the nucleus are shown as bar graphs (mean ± SD; p–r). Significant difference is indicated with asterisks (*n* = 3, *p* < 0.05, Student *t*-test).

In contrast, K-975 modestly decreased the conventional SMAD2/3 target gene *SERPINE1* and had no statistically significant effect on *SMAD7* **(Fig. 2f, g)**. *SMAD2* expression was not altered by either TGF-β1 or K-975 **(Fig. 2h)** and TGF-β1-mediated downregulation of *SMAD3* was not reversed by K-975 **(Fig. 2i)**. K-975 also moderately suppressed phosphorylation of SMAD2/3 in fibroblasts exposed to TGF-β1 **(Fig. 2j)**. Although K-975 primarily inhibits YAP/TAZ interaction with TEAD transcription factors, it also reduced YAP/TAZ nuclear localization **(Fig. 2k–m)**. Nuclear localization of SMAD2/3 was also decreased in K-975-treated cells, but remained high compared to untreated control cells **(Fig. 2k, n)**.

Proximity ligation assay (PLA) demonstrated nuclear interaction between YAP/TAZ-SMAD2/3 and TEAD1-SMAD2/3 increased following TGF-β1 treatment and were reduced by K-975 **(Fig. 2o–r)**, suggesting cooperative interactions between YAP/TAZ, TEAD1, and SMAD2/3 contribute to TGF-β-induced transcription.^17^ Consistently, dual knockdown of YAP and TAZ greatly reduced expression of YAP/TAZ target genes and more weakly reduced expression of SMAD2/3 target genes, similar to K-975 treated cells **(Fig. 3)**.

**Figure 3.**
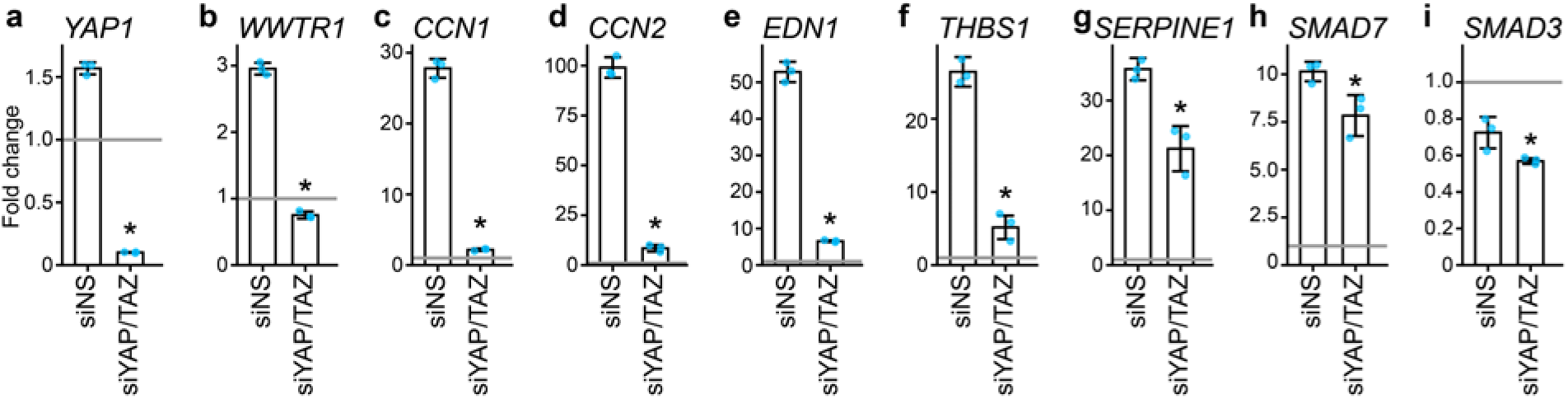
YAP/TAZ gene knockdown decreased expression of both YAP/TAZ and SMAD2/3 target genes in human VF fibroblasts *in vitro* with a modest effect on SMAD2/3. Human VF fibroblasts were treated with nonsense or YAP/TAZ siRNA (siNS or siYAP/TAZ) for two days, and subsequently treated with 10ng/mL TGF-β1 +/- 0.3‒10μM K-975. Gene expression 24 hours after TGF-β1 +/- K-975 treatment was assessed by RT-qPCR (a–i). Fold changes relative to control cells, not with TGF-β1 or K-975, are shown as bar graphs (mean ± SD). Significant difference is indicated as asterisks (*n* = 3, *p* < 0.05, Student’s *t*-test).

Collectively, these data suggest YAP/TAZ and SMAD2/3 both independently and cooperatively regulate TGF-β-responsive transcription, with YAP/TAZ activation playing a more impactful role in driving fibroblast activation. Based on these findings, we hypothesized YAP/TAZ inhibition effectively prevents fibrosis by selectively suppressing pro-fibrotic mechanisms within TGF-β signaling.

### RNA-seq predicted YAP/TAZ to strongly impact TEAD-related transcription compared to SMAD3 target genes

Transcriptomic profiling of VF fibroblasts treated with TGF-β1 identified significant induction of numerous genes including those validated by RT-qPCR **(Fig. 4a, Supplementary Materials Tables S1 and S2)**. Among the most significantly induced genes were *PTHLH*, *PMEPA1*, *BHLHE40*, *PRG4*, *ELN*, and *ESM1*. Several of these genes have been implicated in negative regulation of TGF-β signaling and fibrosis. For example, *PTHLH* both positively and negatively regulates SMAD2/3 depending on exposure time to TGF-β1, *PMEPA1* suppresses SMAD3 phosphorylation, and *PRG4* inhibits SMAD2 phosphorylation and myofibroblast differentiation.^28–31^ Genes induced by TGF-β1 were generally downregulated in response to K-975, with the exception of *PTHLH* and *PMEPA1* **(Fig. 4b)**. Similarly, K-975 had only a modest effect on SMAD2/3 target genes *SERPINE1* and *SMAD7*. In contrast, K-975 significantly reduced YAP/TAZ target genes *CCN1*, *CCN2*, and *THBS1*.

**Figure 4.**
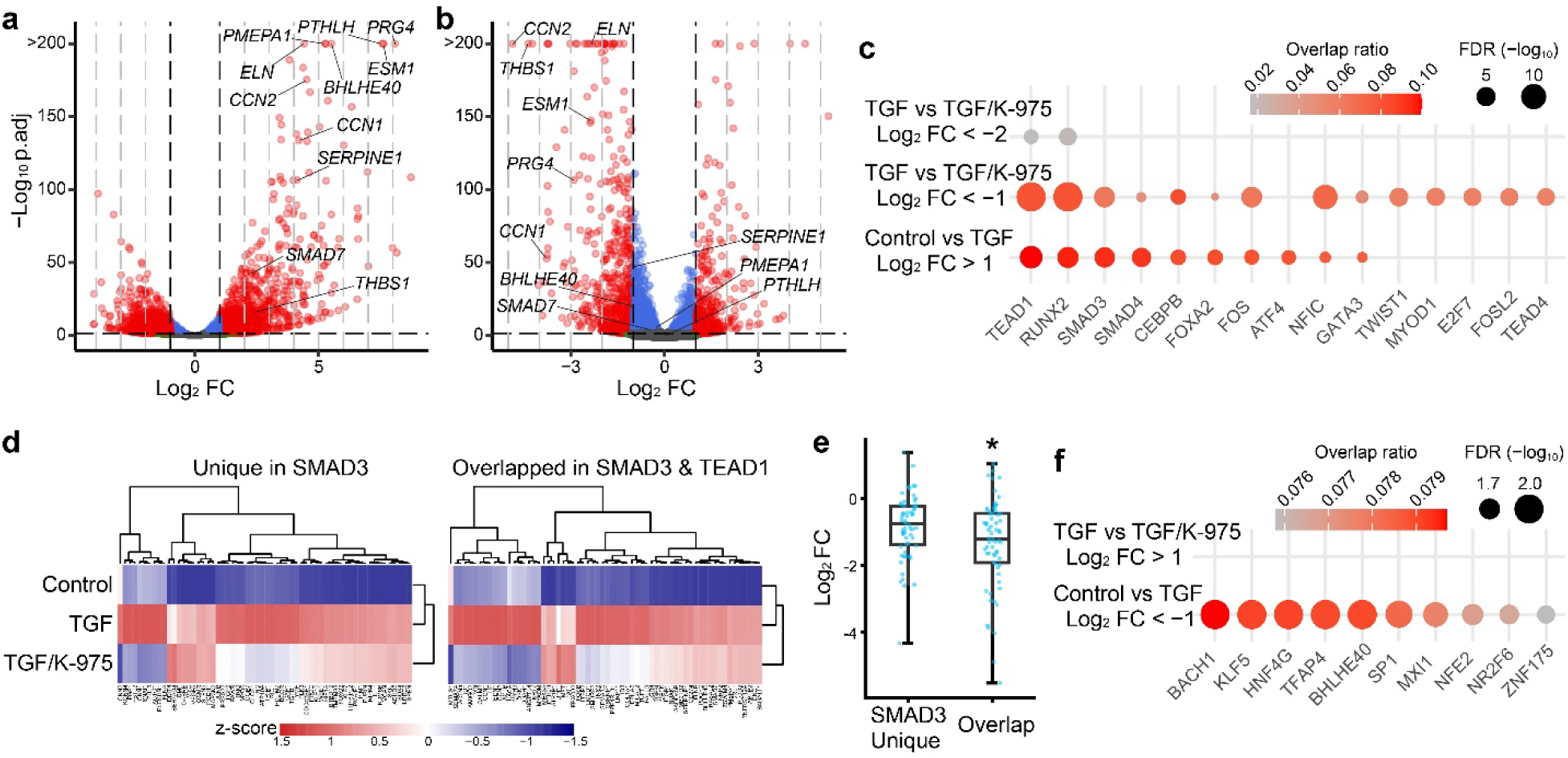
Transcriptome analysis predicted differential impact of YAP/TAZ on SMAD2/3-mediated and non-SMAD2/3 signaling triggered by TGF-β in human VF fibroblasts. Transcriptome of human VF fibroblasts treated with 10ng/mL TGF-β1 +/- 10μM K-975 for 12 h and untreated control cells was assessed by RNA-seq. Volcano plots illustrate relative transcription levels in fibroblasts treated with TGF-β1 alone compared with control cells (a), and in cells treated with TGF-β1 + K-975 compared with cells treated with TGF-β1 alone (b). Red plots indicate differentially expressed genes (DEGs) determined with adjusted *p*-value < 0.05, and log_2_ fold change (FC): < −1 and > 1. To predict transcription factors activated via TGF-β signaling and suppressed by YAP/TAZ inhibition, gene set enrichment analysis (GSEA) was performed using the ReMap ChIP-seq library (c). Top 10 transcription factors, based on false discovery rates (FDR), for genes upregulated by TGF-β1 (log_2_ FC: > 1) and downregulated by K-975 (log_2_ FC: < −1 and < −2) are shown as dot plots, with removal of redundancy (c; full lists of the GESA results are available in Supplementary Materials Tables S3–5). Overlap ratios in the reference gene set and FDRs are indicated by color changes and dot sizes, respectively. Heatmaps illustrate trends of K-975 effect on genes upregulated by TGF-β1 for gene sets that were unique in the SMAD3 reference set and overlapped between the SMAD3 and TEAD1 reference set (d). Changes in transcription levels by K-975 treatment were assessed for the two gene sets and shown as a box-whisker plot (e). Significant difference is indicated as asterisks (*p* < 0.05, Kruskal-Wallis test). SMAD3-unique genes: *n* = 60. Overlapped genes: *n* = 64. GSEA was performed for genes downregulated by TGF-β1 (log_2_ FC: < −1) and upregulated by K-975 (log_2_ FC: > 1) using the ReMap ChIP-seq library, and top 10 transcription factors, based on false discovery rates (FDR), are shown as dot plots, with removal of redundancy (f; a full list of the GSEA results are available in Supplementary Material Tables S6).

To further define transcriptional networks regulated by TGF-β1 and YAP/TAZ inhibition, we performed gene set enrichment analysis (GSEA) using the ReMap database, which integrates chromatin immunoprecipitation sequencing (ChIP-seq)-derived transcription factor data.^32^ Analysis of genes upregulated by TGF-β1 (log_2_ fold change: > 1; adjusted p-value: < 0.05) predicted activation of multiple transcription factors associated with fibrosis and SMAD signaling including SMAD3, TEAD1, RUNX2, FOS, and CEBPB **(Fig. 4c, Supplementary Material Table S3)**.^33–35^ FOXA2, reported to be anti-fibrotic, was also predicted to be activated by TGF-β1.^36^ Many of these transcriptional programs overlapped with genes suppressed by K-975 treatment (log_2_ fold change: < −1; adjusted p-value: < 0.05; **Fig. 4c, Supplementary Material table S4)**.

We subsequently focused on genes more highly impacted by K-975 (log_2_ fold change: < −2; adjusted p-value: < 0.05; **Fig. 4c, Supplementary Material Table S5)**. When the analysis was restricted to genes strongly downregulated by K-975, TEAD1 and RUNX2 remained significantly enriched, whereas SMAD3 was no longer significant **(Fig. 4c, Supplementary Material Table S3)**. To better distinguish YAP/TAZ-dependent from YAP/TAZ-independent components of SMAD3-mediated transcription, SMAD3 reference genes were divided into two categories: 1) SMAD3 unique genes, which are found in the SMAD3 reference gene set, but not in the TEAD1 gene set, and 2) TEAD1/SMAD3-overlapping genes, which are found in both SMAD3 and TEAD1 reference gene sets. Genes upregulated by TGF-β1 were more frequently suppressed by K-975 treatment, returning to near or below basal levels in the TEAD1/SMAD3-overlapping genes. SMAD3-unique genes were less affected **(Fig. 4d, e)**. These findings suggest YAP/TAZ inhibition disrupts TEAD-associated, profibrotic transcriptional programs that cooperate with SMAD3 signaling.

Genes downregulated by TGF-β1 treatment (log_2_ fold change: < −1; adjusted p-value: < 0.05) were enriched in multiple transcription factors known to be involved in cancer progression, the response to stress, and the immune system; negative regulation of these transcription factors by TGF-β signaling was predicted **(Fig. 4f, Supplementary Material Table S6)**. Several of these transcription factors, such as BACH1, KLF5, and SP1, reportedly contribute to fibrosis and/or fibroblast activation.^37–39^ However, no significantly enriched transcription factors were identified among genes upregulated by K-975 treatment (log_2_ fold change: > 1; adjusted p-value: < 0.05). These findings suggest YAP/TAZ primarily targets pro-fibrotic transcriptional pathways activated by TGF-β, but have relatively little impact on transcriptional programs suppressed by TGF-β signaling.

### ChIP-seq identified distinct chromatin occupancy between SMAD2/3 and YAP/TAZ

Since reference gene sets were derived from non-VF cell types, we performed ChIP-seq to directly examine genomic occupancy profiles of SMAD2/3 and YAP/TAZ in VF fibroblasts following TGF-β stimulation. In untreated cells, ChIP-seq peaks identified for SMAD2/3, YAP, and TAZ had different profiles **(Fig. 5a)**. TGF-β1 treatment resulted in some peak overlap between SMAD2/3, YAP, and TAZ, but ∼67% of peaks for SMAD2/3 did not overlap with YAP/TAZ and ∼80% of peaks for YAP/TAZ did not overlap with SMAD2/3 **(Fig. 5a, b)**, indicating these transcriptional regulators largely occupy separate genomic regions.

**Figure 5.**
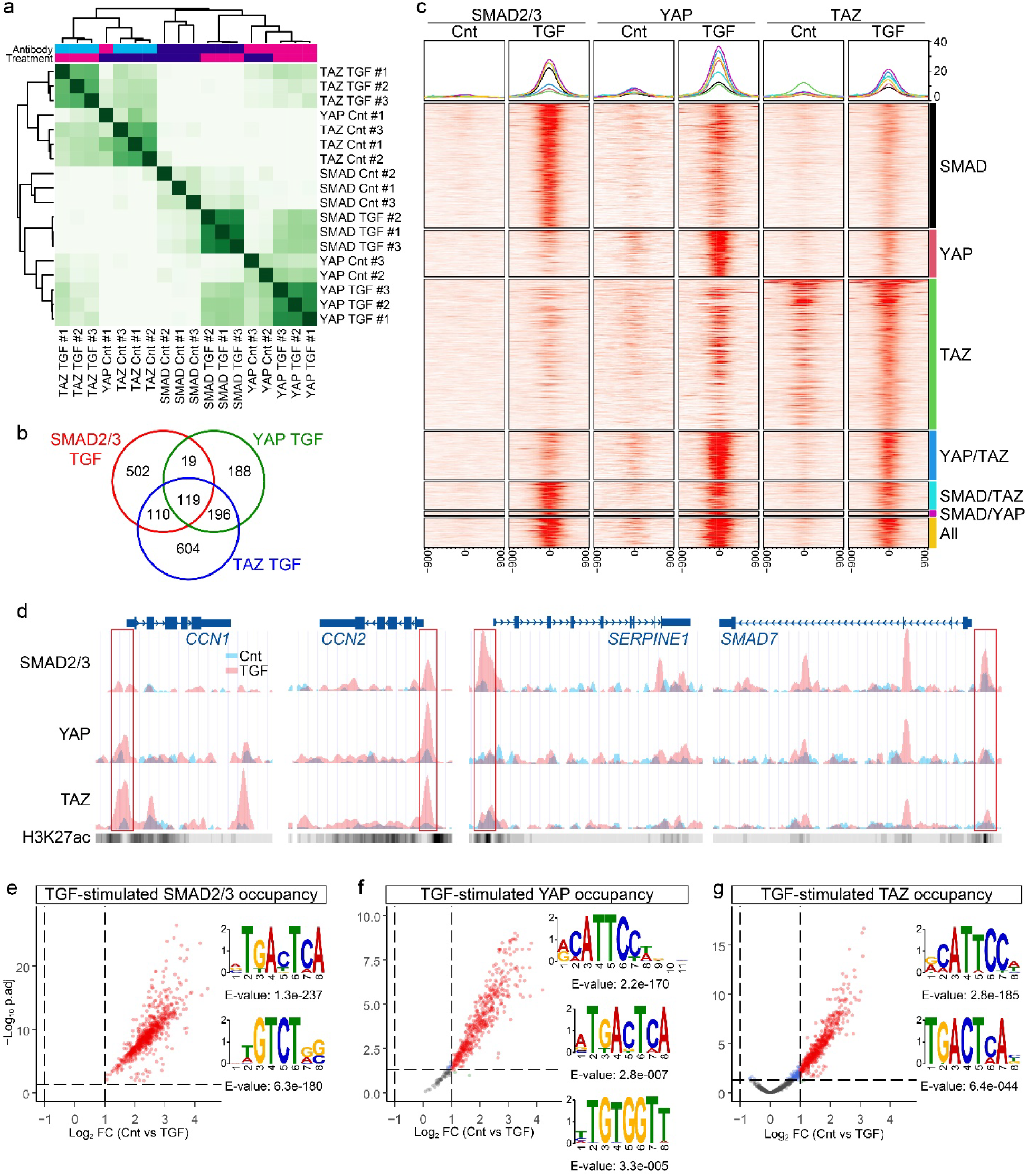
ChIP-seq predicted direct cooperation between YAP/TAZ and SMAD2/3 is not a major contributor to TGF-β signaling. DNA occupancy of SMAD2/3 and YAP/TAZ in Human VF fibroblasts treated with or without 10ng/mL TGF-β1 were assessed by ChIP-seq. A correlation heatmap, created based on read counts, visualizes relationships across samples derived from SMAD2/3, YAP, and TAZ antibody binding (a). Overlap between SMAD2/3, YAP, and TAZ peak sites are illustrated as a Venn diagram (b). Profiles of ChIP-seq peaks are shown as a profile plot and complex heatmap (c). Peak sets of unique binding sites for SMAD, YAP, and TAZ, and overlapped sites for two or three of these proteins were assessed, and results from different peak sets are shown as distinct colors in the profile plot. Chromatin occupancy of SMAD2/3, YAP, and TAZ around coding regions of *CCN1*, *CCN2*, *SERPINE1*, and *SMAD7* are visualized (d). DNA occupancy stimulated by TGF-β1 treatment was determined based on increased read counts at ChIP-seq peaks (log_2_ fold change (FC) > 1; adjusted *p*-value < 0.05), and motif discovery was performed (e–g). Volcano plots show read counts in TGF-β1-treated cells compared with control cells. Discovered DNA motifs are shown beside the volcano plots.

TGF-β1 increased chromatin occupancy of SMAD2/3, YAP, and TAZ at their peak sites. The impact on TAZ occupancy, however, was modest **(Fig. 5c)**. At established target genes, TGF-β1 increased occupancy of YAP/TAZ upstream transcription start sites of known YAP/TAZ target genes *CCN1* and *CCN2*, consistent with our transcriptional data **(Fig. 5d)**. However, a clear increase in occupancy was only observed for SMAD2/3 upstream transcription start sites of *SERPINE1* and *SMAD7*. To focus specifically on TGF-β-responsive occupancy, we analyzed peaks significantly enriched following TGF-β1 stimulation (log_2_ fold change: > 1; adjusted p-value: < 0.05; **Fig. 5e–g**). Genome loci identified for TGF-β1-stimulated occupancy of SMAD2/3, YAP, and TAZ are listed in **Supplementary Materials Tables S7–9**.

Motif discovery analysis using the MEME algorithm^40^ revealed distinct DNA-occupancy signatures for SMAD2/3 and YAP/TAZ. TGF-stimulated occupancy of SMAD2/3 was enriched at TGAXTCA and GTCT motifs, corresponding to AP-1 (FOS/JUN) and SMAD binding motifs **(Fig. 5e)**.^41,42^ The TGAXTCA motif was also identified for TGF-stimulated occupancy of YAP and TAZ. However, the SMAD motifs (GTCT) were not significantly enriched within YAP/TAZ peaks **(Fig. 5f, g)**. Instead, YAP/TAZ occupancy was associated with CATTCCA and TGTGGT motifs, corresponding to TEAD and RUNX binding sites.^43,44^ These findings indicate direct cooperation of SMAD2/3 and YAP/TAZ for binding to gene regulatory elements is likely limited. However, complementary motif analysis using the CentriMo database identified SMAD and TEAD binding motifs as targets of both YAP/TAZ and SMAD2/3 **(Supplementary Materials Tables S10–12)**,^45^ supporting the possibility of cooperative interactions between SMAD2/3, YAP/TAZ, and TEAD transcription factors at select regulatory elements in VF.

### Multi-omic analysis predicted gene sets activated by SMAD2/3 and YAP/TAZ and differential impact of YAP/TAZ inhibition

To predict transcriptional programs activated by chromatin occupancy of SMAD2/3 and YAP/TAZ, TGF-stimulated occupancy identified by ChIP-seq were mapped to promoter- and enhancer-like regions encompassed within the candidate *cis*-regulatory elements (cCREs) database, which encompasses gene regulatory elements identified based on DNase-seq data, H3K4me3 binding, H3K27ac binding, and Hi-C analysis.^46^ **Supplementary Material Table S13** lists promoter/enhancer-like regions identified for TGF-stimulated chromatin occupancy of SMAD2/3 and YAP/TAZ, as well as genes regulated by these regions. Subsequently, genes upregulated by chromatin occupancy of SMAD2/3 and YAP/TAZ under TGF-β stimulus were predicted by cross-referencing datasets of the predicted gene regulation from ChIP-seq data and upregulated genes by TGF-β1 treatment from RNA-seq data (log_2_ fold change: > 1; adjusted p-value: < 0.05; **Supplementary Material Table S14**). These genes were classified into two groups: 1) SMAD2/3-unique targets, exclusively identified for SMAD2/3; and 2) YAP/TAZ-related targets, identified for YAP/TAZ regardless of overlap with SMAD2/3. We explored biological events regulated by these gene sets using the Reactome pathway database.^47^ SMAD2/3-unique target genes were enriched in reference gene sets related to cell-ECM interaction and TGF-β and SMAD2/3/4 signaling, including downregulation of SMAD2/3/4 **(Fig. 6a, Supplementary Material Table S15)**. YAP/TAZ-related target genes were likely associated with broader signaling events related to fibroblast activation such as Rho GTPase, ECM organization, and non-integrin-based ECM interaction **(Supplementary Material Table S16)**.^48^ In addition, RAS/MAPK and multiple receptor tyrosine kinase signaling (c-MET, FGFRs, and IGFR), which broadly induce MAPK signaling, were predicted for YAP/TAZ-related target genes, implying YAP/TAZ interaction with non-canonical TGF-β signaling via MAPKs.^8,49^ Despite putative interactions with various signaling pathways, including activation of TGF-β and SMAD2/3 signaling, YAP/TAZ-related target genes were relatively poorly enriched in the gene set related to downregulation of SMAD2/3/4.

**Figure 6.**
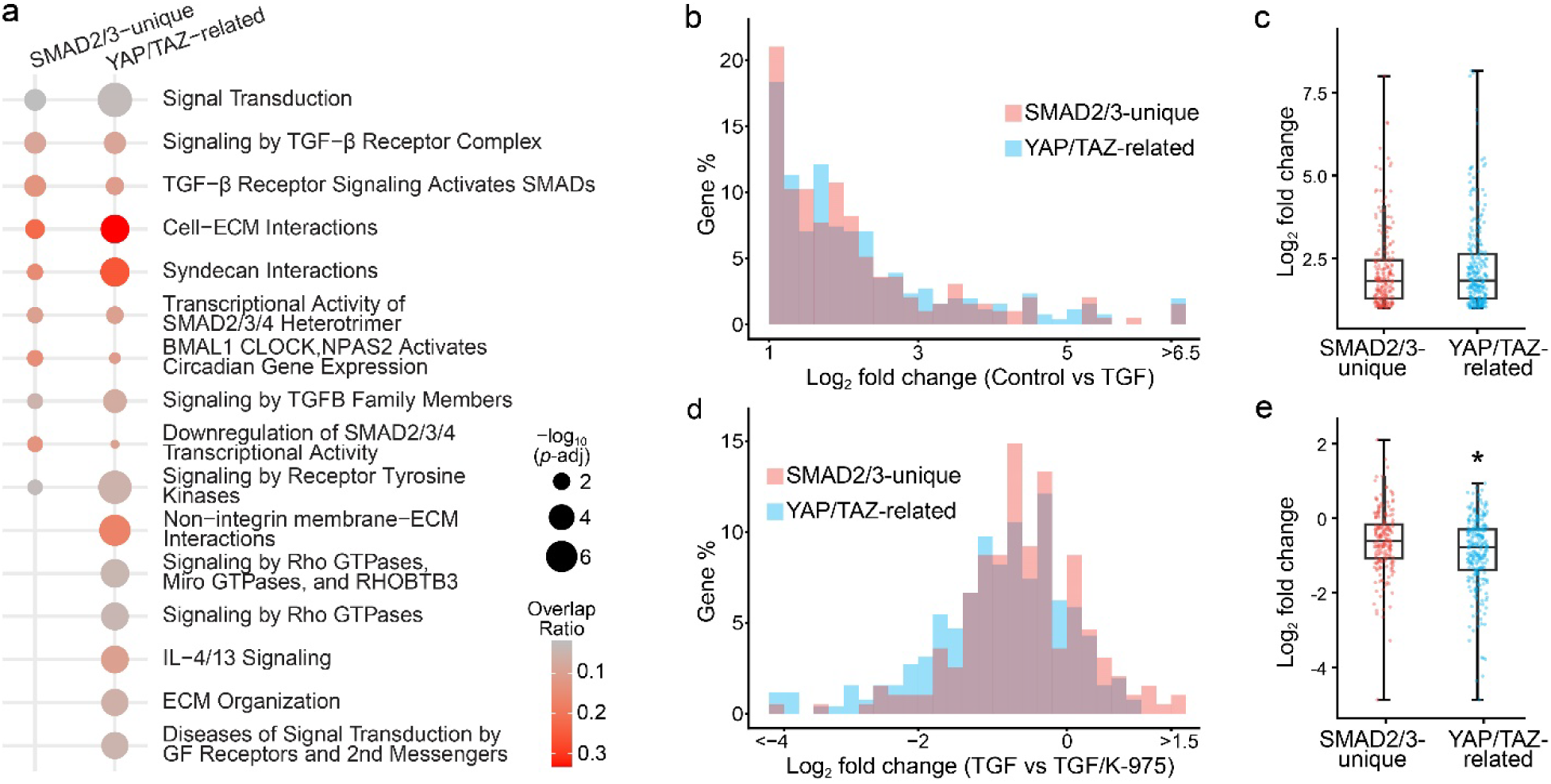
SMAD2/3-unique and YAP/TAZ-related target genes differentially impacted signaling pathways and sensitivity to K-975 treatment in VF fibroblasts. SMAD-unique and YAP/TAZ-related target genes were determined by multi-omic analyses with the RNA-seq and ChIP-seq data. Gene set enrichment analysis was performed with the two datasets using the Reactome 2024 library (a). Top 10 terms (based on adjusted *p*-value) identified for respective datasets are shown as dot plots, with removal of redundancy (a full list of the GESA results are available in Supplementary Material Table S16). Expression levels in TGF-β1-treated cells relative to control cells, determined by RNA-seq, are plotted as histogram and box-whisker plots for the SMAD2/3-unique and YAP/TAZ-related target genes (b, c). Expression levels for TGF-β1/K-975 combination treatment, relative to TGF-β1 treatment, are also plotted (d, e). Significance is indicated as an asterisk (*p* < 0.05, Kruskal-Wallis test). SMAD2/3-unique targets, *n* = 198. YAP/TAZ-related targets, *n* =256.

We next examined the impact of TGF-β and YAP/TAZ inhibition on transcription of SMAD2/3-unique and YAP/TAZ-related gene sets. Both SMAD2/3-unique and YAP/TAZ-related targets were induced to a similar extent by TGF-β1 treatment, with no significant difference in the magnitude of transcriptional activation between groups **(Fig. 6b, c)**. In contrast, YAP/TAZ-related target genes trended toward suppression by K-975 treatment **(Fig. 6d)**. K-975-mediated downregulation was greater in YAP/TAZ-related target genes compared to SMAD2/3-unique target genes **(Fig. 6e)**.

Collectively, our *in vitro* findings suggest YAP/TAZ promote fibroblast activation through multiple pro-fibrotic signaling pathways downstream of TGF-β signaling, while exerting only limited effects on the negative feedback mechanisms associated with canonical SMAD2/3 signaling. These data further support the therapeutic potential for YAP/TAZ inhibition for VF fibrosis.

### Pharmacological YAP/TAZ inhibition prevents VF fibrosis in a rat iatrogenic injury model

To evaluate the anti-fibrotic effect of YAP/TAZ inhibition *in vivo*, K-975 was administered to injured VFs in a rat iatrogenic VF injury model seven days after VF injury, corresponding to the subacute phase of wound healing. Immunofluorescence analysis performed one day after treatment demonstrated that both 3μM and 10μM K-975 reduced nuclear localization of YAP and TAZ **(Fig. 7a–c)**. In contrast, inhibition of SMAD2/3 and phosphorylated SMAD2/3 nuclear localization was observed only with 3μM K-975 treatment **(Fig. 7a, d, e)**. Moderate YAP/TAZ inhibition by 3μM K-975 may be more efficient to suppress both YAP/TAZ and SMAD2/3 signaling under the complexity of the inflammatory milieu with multiple cell types and signaling events.

**Figure 7.**
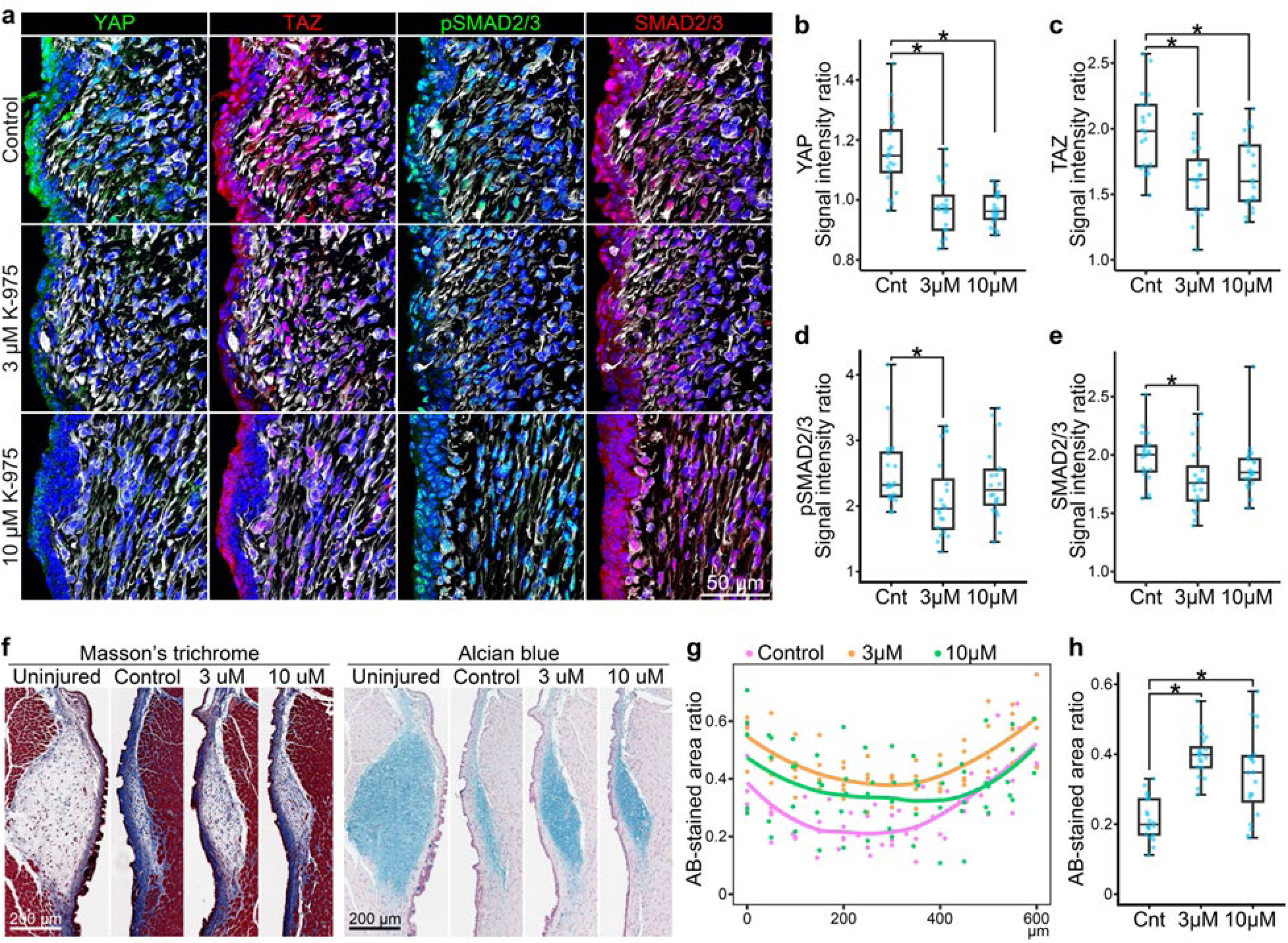
K-975 inhibited post-injury YAP/TAZ nuclear localization and fibrosis in the rat VF. Vehicle, 3µM K-975, or 10µM K-975 was injected into the injured sites in the rat VF iatrogenic injury model seven days after surgery. Immunofluorescence images of coronal cryosections show distributions of YAP (green), TAZ (red), phosphorylated SMAD2/3 (pSMAD2/3; green), and total SMAD2/3 (red) (a). Nuclei and mesenchymal cells were counterstained using DAPI (blue) and anti-vimentin antibody (grey). The intensity of immunofluorescence signal in the DAPI-stained region, relative to the vimentin-positive region, were measured in four sections per rat (b–e). The sections were obtained at 90–410µm from the anterior end of the arytenoid cartilage with at least 90µm intervals between sections. Statistical significance is denoted as asterisks (*p* < 0.05, Kruskal-Wallis and post-hoc Wilcoxon rank sum tests with Bonferroni adjustment, *n* = 20). Coronal paraffin-embedded sections were stained by Masson’s trichrome and Alcian blue staining (f). Alcian blue (AB)-stained areas in VF lamina propria were compared between injured and uninjured VFs on each section, and ratios of the stained areas are plotted (g, h). Data obtained within 0–600µm from the anterior end of the arytenoid cartilage (g). Data from four sections per rat, obtained at 85–415µm with at least 90µm intervals between sections (h). Statistical significance is denoted as asterisks (*p* < 0.05, Kruskal-Wallis and post-hoc Wilcoxon rank sum tests with Bonferroni adjustment, *n* = 20).

The effect of K-975 on VF fibrosis was assessed histologically 56 days after injury. This timepoint was selected to reflect scar maturity. Masson’s trichrome staining, which detects collagen deposition and fibrosis, demonstrated 3μM and 10μM K-975 reduced fibrotic tissue compared to untreated injured controls **(Fig. 7f)**. Alcian blue staining, which marks glycosaminoglycan-rich ECM, was maintained outside the fibrotic lesion. Alcian blue positivity significantly increased in response to 3μM and 10μM K-975, confirming anti-fibrotic effects of YAP/TAZ inhibition *in vivo* **(Fig. 7f–h)**. These findings confirm pharmacological inhibition of YAP/TAZ attenuates VF fibrosis and promote preservation of ECM composition *in vivo*.

## DISCUSSION

Given the anatomical position of the VFs and their involvement in multiple critical biological processes, VF fibrosis presents a significant clinical challenge with substantial impact on quality of life. The lack of effective therapeutic approaches for VF fibrosis also limits surgical options for other VF lesions, due to the risk of iatrogenic injury resulting in exacerbation of the fibrotic response. The role of TGF-β signaling in fibrosis has been extensively reported. However, signaling molecules other than SMAD2/3 have emerged as therapeutic targets because of their unique roles in the fibrotic response. The current study provides insight into the differential impact of YAP/TAZ across TGF-β-induced transcription. Our findings suggest YAP/TAZ inhibition is a promising approach to suppress the fibrotic response via selective suppression of the pro-fibrotic component of TGF-β signaling.

YAP/TAZ, as a core signaling molecule in the Hippo signaling pathway, was originally investigated in the context of organ growth and development.^50,51^ More recently, due to interactions with multiple signaling molecules associated with fibroblast activation such as Rho GTPase, β-catenin, and glucocorticoid receptor, YAP/TAZ has emerged as an intriguing therapeutic target for fibrosis.^48,52–55^ TGF-β drives both SMAD2/3 and YAP/TAZ activation, and cooperation of these molecules (including physical interaction) is thought to be an important mechanism of fibroblast activation.^56^ However, our gene expression data suggest limited impact of YAP/TAZ inhibition on SMAD2/3 target genes (e.g., SMAD7) compared to YAP/TAZ target genes (e.g., CCN2), despite significant suppression of marker genes of fibroblast activation. Our transcriptomic analysis predicted YAP/TAZ only partially mediated SMAD2/3-dependent transcription and did not specifically influence transcription factors negatively regulated by TGF-β signaling.

Our ChIP-seq data showing induction by TGF-β1 to distinct genomic loci support substantially non-overlapping roles of YAP/TAZ and SMAD2/3. Approximately two-thirds of SMAD2/3 genomic occupancy sites were not shared with YAP/TAZ. The conventional binding element for SMAD2/3 was not strongly associated with YAP/TAZ recruitment. Instead, JUN/FOS and RUNX binding motifs, in addition to the TEAD binding motif, were identified as target sites of YAP/TAZ. Activator protein-1 (AP-1), a dimeric transcription factor composed of FOS and JUN family molecules, physically interacts with SMAD2/3 and drives gene expression.^57^ Close localization of AP-1 and SMAD binding motifs are frequently found in upstream target genes such as *SERPINE1* and *SMAD7*, and cooperation of AP-1 and SMAD2/3 is required for efficient gene expression.^58,59^ However, our ChIP-seq data suggest YAP/TAZ occupancy at the AP-1 binding motif was mainly mediated via SMAD2/3-independent mechanisms, since SMAD2/3 and YAP/TAZ chromatin occupancy were largely distinct. AP-1 can be activated through non-canonical TGF-β signaling such as activation of MAPKs and Rho GTPases; AP-1 dependency has also been described for efficient induction of YAP/TAZ target genes.^8,60,61^ In addition, AP-1 and YAP/TAZ physically interact and co-occupy chromatin to synergistically activate genes related to cancer growth.^62,63^ Further studies investigating SMAD2/3-independent chromatin co-occupancy of AP-1 and YAP/TAZ are warranted to clarify YAP/TAZ roles in the non-canonical response to TGF-β signaling in VF fibroblasts.

Recent reports suggest RUNX1/2 serves as another fibrotic signaling factor induced by TGF-β. RUNX1 was activated by TGF-β1 in pulmonary fibroblasts.^64^ RUNX1 gene knockdown and knockout inhibited dermal and cardiac fibroblast activation.^65,66^ Overexpression of RUNX2 exacerbated hepatic fibrosis.^33^ In addition, cooperation between the YAP-TEAD complex and RUNX2 in cardiac fibroblast activation has been suggested.^67^ These data, combined with our current findings, suggest cooperation of YAP and RUNX1/2 may be another regulatory mechanism underlying TGF-β-induced fibroblast activation.

Multiomic analyses with RNA-seq and ChIP-seq datasets indicated transcription mediated by chromatin occupancy of SMAD2/3 and YAP/TAZ differentially influenced cell signaling pathways induced by TGF-β. In contrast to SMAD2/3-unique target genes, which primarily regulate canonical TGF-β/SMAD signaling, YAP/TAZ-related target genes were associated with a number of distinct signaling pathways. Signaling pathways related to ECM organization, cell-ECM interaction (via both integrin and non-integrin molecules), MAPKs, and Rho GTPases were associated with YAP/TAZ-related target genes. These events are crucial for synthesis and crosslinking of ECM and tissue contraction via cytoskeletal rearrangement, indicating a role for YAP/TAZ in altered tissue architecture and stiffness. Of note, YAP/TAZ and Rho GTPases regulate mechanosensitive signal transduction and are activated by a stiff matrix.^68,69^ Activation of Rho/ROCK signaling drives contractile forces via stress fiber formation.^70,71^ Cytoskeletal rearrangement driven by Rho/ROCK signaling induces YAP/TAZ nuclear localization via release of YAP/TAZ from actin filaments.^69^ IL-4 and IL-13 were also associated with YAP/TAZ-related target genes. These Th2 cytokines reportedly activate pulmonary fibroblasts.^72^

With regard to translational application, our *in vitro* findings indicate pharmacological YAP/TAZ inhibition preferentially suppressed YAP/TAZ-mediated transcription, including expression of target genes like *CCN1* and *CCN2* with key roles in fibrosis more than SMAD2/3-unique transcription. Selective suppression of pro-fibrotic programs within TGF-β signaling supports therapeutic potential of a YAP/TAZ-inhibiting strategy for fibrosis. The anti-fibrotic effects of YAP/TAZ inhibition were confirmed in our rat VF fibrosis model. Our *in vivo* findings suggest a single injection of K-975, a YAP/TAZ inhibitor, prevents VF fibrosis. However effective concentrations of K-975 differed between *in vitro* and *in vivo* experiments. A lower concentration (3µM) of K-975 was sufficient to prevent VF fibrosis *in vivo,* but less effectively reduced fibrotic gene expression *in vitro* compared to 10µM. In addition, 10µM K-975 did not significantly inhibit nuclear localization of SMAD2/3 *in vivo*. The effects of YAP/TAZ on non-fibroblast cells may be the source of the discrepancy between effective concentrations *in vitro* and *in vivo*. For example, YAP gene knockout promotes macrophage polarization toward the M2 phenotype, a pro-fibrotic phenotype secreting TGF-β.^73,74^ Further studies are warranted to clarify potential side effects of YAP/TAZ inhibition on non-fibroblast cells in the VF.

Collectively, the current study provides key insights regarding YAP/TAZ as a potential therapeutic target for VF fibrosis with significant implications for fibrotic processes beyond the upper aerodigestive tract. Our findings indicate YAP/TAZ mediates the fibrotic response with only a modest impact on negative feedback mechanisms of TGF-β/SMAD signaling.

## METHODS

### Cells

An immortalized human VF fibroblast line (HVOX), created by and maintained in our laboratory, was expanded to P16‒17 using Dulbecco’s Modified Eagle’s Medium (DMEM) containing 10% fetal bovine serum (FBS) and 1% antibiotic/antimycotic (Life Technologies, Grand Island, NY) at 37°C, 5% CO_2_. Prior to treatment with 10ng/mL TGF-β1 and 0.3‒10μM K-975 (MedChemExpress, Monmouth Junction, NJ, USA), cells were rinsed twice with phosphate-buffered saline (PBS, pH=7.4) and exposed to serum-free DMEM for 24 h. For double gene knockdown of YAP1 and WWTR1 (encoding YAP and TAZ), cells were treated with Silencer Select Pre-designed siYAP1 (sense: GGUGAUACUAUCAACCAAAtt; antisense: UUUGGUUGAUAGUAUCACCtg) and siWWTR1 (sense: GUACUUCCUCAAUCACAUAtt; antisense: UAUGUGAUUGAGGAAGUACct) for two days before TGF-β1 treatment. Nonsense siRNA was used for the control.

### Animals

Following prior approval by the Institutional Animal Care and Use Committee (IACUC) at the New York University Grossman School of Medicine, all animal experiments were performed in accordance with relevant institutional guidelines and regulations (IACUC protocol ID: PROTO202100053). Male Sprague–Dawley rats were obtained from Charles River Laboratories (Hopkinton, MA, USA). VF injury and K-975 injection were performed at age 14 after > 4 days of acclimation, as detailed below. Five rats were employed for three conditions (control, 3μM and 10μM K-975) and two time points for tissue harvest (one day and seven weeks after injection); thirty rats were employed in total. At the time points of tissue harvest, all animals were euthanized by exposure to CO_2_ gas.

### Rat VF injury and K-975 injection

Intraperitoneal injection of atropine sulfate (50µg/kg) and topical administration of 1% lidocaine and 0.5mg/mL epinephrine was performed for rats under paralytic anesthesia induced by intraperitoneal injection of ketamine hydrochloride (90mg/kg) and xylazine hydrochloride (8mg/kg). The right VF mucosa was incised using a 25-gauge needle under endoscopic guidance (Visera Elite, Olympus, Tokyo, Japan). PBS, 3µM K-975 and 10µM K-975 (containing 1% dimethylsulfoxide as a vehicle) were injected via a transoral approach (5µL) into the injured site seven days after injury. Larynges were harvested one day and seven weeks after injection for immunofluorescence and histological examinations.

### Quantitative real-time polymerase chain reaction (RT-qPCR)

Cells were treated with or without 0.3‒10μM K-975 for 24 hours. Following RNA extraction and reverse transcription using the RNeasy Mini Kit (Qiagen, Valencia, CA) and High-Capacity cDNA Reverse Transcription Kit (Applied Biosystems), real-time polymerase chain reaction was performed using the TaqMan Gene Expression kit (Life Technologies) and Taqman primer probes, listed in **Supplementary Material Table S17**. Amplification signals were detected using StepOne Plus (Applied Biosystems). Expression levels relative to GAPDH were quantified by the ΔΔCt method.

### Western blotting

Cells were treated with or without 1‒10μM K-975 for 12 hours and 24 hours and collected using PBS supplemented with 0.1% Triton X-100, Halt Protease Inhibitor Cocktail (Thermo Scientific), Halt Phosphatase Inhibitor Cocktail (Thermo Scientific), 5mM EDTA Solution, Calyculin A (Cell Signaling). Following heat denaturation with 4x Laemmli Sample buffer (Bio-Rad) and 2-mercaptoethanol, proteins in the samples were separated by sodium dodecyl sulfate-polyacrylamide gel electrophoresis and transferred to PVDF membranes (Invitrogen). The membranes were incubated with 5% BSA (Fisher Scientific) at 4°C overnight, followed by incubation with primary and horse radish peroxidase (HRP)-conjugated secondary antibodies **(Supplementary Material Table S18)**. Signals derived from HRP reaction with SuperSignal™ West Femto Maximum Sensitivity Substrate (Pierce Biotechnology, Rockford, IL) were detected using ChemiDoc MP (Bio-Rad Laboratories, Hercules, CA).

### Immunostaining and proximity ligation assay (PLA)

Cells treated with 10ng/mL TGF-β1 +/- 10μM K-975 for 12 hours were fixed with 4% paraformaldehyde. Coronal cryosections of larynges were fixed using 10% formalin. Specimens were permeabilized by exposure to 0.1% Triton X-100. Following incubation with 1% BSA, cells were reacted with primary antibodies, and subsequently with secondary antibodies conjugated with Alexa Fluor or PLA probes **(Supplementary material Table 18)**. For PLA, cells were further treated with ligase and polymerase following manufacturer instructions. Nuclei were counterstained with DAPI. Fluorescent signals were observed using Zeiss700 and Zeiss880 confocal laser microscopes (Zeiss, Oberkochen, Germany). DAPI-stained areas were determined as nuclear regions. For cultured cells, signals of antibody reaction and PLA in nuclear regions were quantified using ImageJ (National Institutes of Health). For tissue sections, vimentin-positive areas outside DAPI-stained areas were determined as cytoplasmic areas, and ratios of antibody signals in nuclear regions against cytoplasmic regions were quantified.

### Histology

Coronal paraffin-embedded sections were obtained using rat larynges fixed with 4% paraformaldehyde. Masson’s trichrome and Alcian blue staining was performed for the observation of collagen and glycosaminoglycans (including hyaluronic acid) in the VF mucosa. Nuclear Fast Red staining was performed to counterstain nuclei in Alcian blue-stained sections. Microscopic images were obtained using an Aperio AT2 slide scanner (Leica Biosystems, Nussloch, Germany). Alcian blue-stained areas were quantified at the left uninjured and right injured/treated VF mucosae on each section. Ratios of Alcian-blue stained areas in the right VF mucosa compared to the left side were calculated to assess preservation of the hyaluronan-rich ECM characteristic of the VF.

### RNA sequencing (RNA-seq)

Cells were treated with or without 10ng/mL TGF-β1 alone or in combination with 10μM K-975 for 12 hours. RNA was isolated using RNA extraction with the RNeasy Mini Kit. Following RNA quality check using the 2100 Bioanalyzer (Agilent Technologies, Santa Clara, CA, USA), RNA-seq libraries were prepared using the Truseq Stranded mRNA kit (Illumina, San Diego,CA, USA). Sequencing was performed in NovaSeq X+ (Illumina). Trimming, quality assessment, and alignment of reads were performed on the Rosalind analysis platform (https://www.rosalind.bio/). The human genome assembly hg38 was employed as a reference for read alignment. DEseq2 was used to calculate fold changes and adjusted *p*-values. Five gene sets were derived from the transcriptome data based on log_2_ fold changes and response to TGF-β1 or K-975: 1) >1 log_2_ fold change by TGF-β1 compared with control; 2) <−1 log_2_ fold change by TGF-β1 compared with control; 3) >1 log_2_ fold change by TGF-β1/K-975 compared with TGF-β1 alone; 4) >2 log_2_ fold change by TGF-β1/K-975 compared with TGF-β1 alone; 5) <−1 log_2_ fold change by TGF-β1/K-975 compared with TGF-β1 alone. GSEA was performed with these gene sets using ChEA3 with the ReMap2022 database.^32^

### Chromatin immunoprecipitation sequencing (ChIP-seq)

Chromatin immunoprecipitation was performed as described previously, with some modifications.^75^ Briefly, cells treated with or without 10ng/mL TGF-β1 for 12 hours were fixed in 1% formaldehyde. Chromatin was isolated using a nuclei isolation buffer containing Igepal, with support of brief sonication (6 cycles of 5-sec sonication at a middle power) using Bioruptor Pico (Diagenode, Seraing, Belgium).^76^ Following replacement of the nuclei isolation buffer with the RIPA buffer, DNA was sheared by 11 cycles of 30-sec sonication at a high power. Isolated chromatin was incubated with antibody-bound Dynabeads protein A (Invitrogen, Carlsbad, CA, USA) at 4°C. After washing the magnetic beads and RNase A treatment, DNA fragments bound to the beads were eluted by heat denaturation in combination with proteinase K treatment and reverse crosslinking using 300mM NaCl. Input controls were prepared from sheared DNA samples by skipping incubation with magnetic beads. DNA fragments were isolated using Zymo ChIP DNA Clean & Concentrator (Zymo Research, Irvine, CA, USA). DNA libraries were prepared using NEBNext Ultra II DNA Library Prep Kit (New England biolabs, Ipswich, MA, USA) and sequenced in NovaSeq X+. Trimming, quality assessment, and alignment of reads were performed on the Rosalind analysis platform. The human genome assembly hg38 was employed as a reference for read alignment. Unmapped and low-quality reads, PCR duplicates, and blacklist regions were removed using samtools and bedtools. Peaks were identified using MACS2. Diffbind was employed to analyze profiles of peaks and create heatmaps. Peaks identified in at least two replicates were determined as consensus peaks and further analyzed to identify overlapping peaks between conditions. Normalized reads were compared between control and TGF-treated cells at sites of consensus peaks identified for TGF-treated cells. TGF-stimulated occupancy was determined based on increased normalized reads via DEseq2 analysis (1 > log_2_ fold change, 0.05 < adjusted p-value). To visualize peaks at specific genome loci on UCSC genome browser (https://genome.ucsc.edu/), mapped reads of replicates were merged using samtools after normalization of read depths, and converted into bigWig files using deeptools. For motif discovery and known motif analysis, 700 bp sequences at TGF-stimulated occupancy sites were analyzed using MEME-ChIP (https://meme-suite.org/).

### Multiomics with RNA-seq and ChIP-seq datasets

TGF-stimulated occupancy sites determined by ChIP-seq were located for promoter-like sites (PLSs) and proximal and distal enhancer-like sites (pELSs and dELSs) based on the human candidate *cis*-regulatory element (cCRE) dataset available at SCREEN (https://screen.wenglab.org/).^46^ To predict gene regulation by TGF-stimulated chromatin occupancy of SMAD2/3 and YAP/TAZ, protein-coding genes that are closest from PLSs and linked to ELSs via 3-dimensional chromatin interaction (based on Hi-C data) were identified using the same database. Genes upregulated by TGF-stimulated chromatin occupancy of SMAD2/3 and YAP/TAZ were predicted via cross-referencing datasets of the predicted gene regulation on ChIP-seq basis and upregulated genes by TGF-β1 treatment on RNA-seq basis (log_2_ fold change: > 1; adjusted p-value: < 0.05). These genes were classified into two gene sets: 1) SMAD2/3-unique targets, exclusively identified for SMAD2/3; and 2) YAP/TAZ-related targets, identified for YAP/TAZ regardless of overlapping with SMAD2/3. Signaling pathways activated by these gene sets were predicted by GSEA on the Enrichr analysis platform (https://maayanlab.cloud/Enrichr/) using Reactome Pathway 2024 as a reference, with the threshold of adjusted p-value < 0.05.^47,77^ In addition, levels of gene upregulation by TGF-β and downregulation by K-975 were assessed for these gene sets by referring to fold changes of normalized transcription levels in the RNA-seq data.

### Statistical Analyses

*In vitro*, three replicates were employed for qPCR, RNA-seq, and ChIP-seq, and five replicates were employed for immunofluorescence and PLA. Student’s t-test was performed for comparison of two conditions in qPCR, immunofluorescence, and PLA data. One-way analysis of variance and Dunnet’s test were performed to compare effects of TGF-β1 alone and combinational treatment of TGF-β1 and varied concentrations of K-975 in qPCR data. *P* < 0.05 was considered significant. Omics data analysis is detailed in the corresponding sections.

*In vivo*, five animals were employed for each condition. Data were obtained from four sections with at least 90µm intervals between sections, within ranges 90–410µm and 85–415µm from the anterior end of the arytenoid cartilage for fluorescently labeled and Alcian blue-stained sections, respectively. Kruskal-Wallis and post-hoc Wilcoxon rank sum tests with Bonferroni adjustment were performed to test significance of the effects of 3µM and 10µM K-975 compared to the control group. *P* < 0.05 was considered significant.

## Acknowledgements

We gratefully acknowledge the financial support provided by the National Institutes of Health/National Institute on Deafness and Other Communication Disorders (R21 DC020993). We thank NYU Langone’s Experimental Pathology Research Laboratory (RRID:SCR_017928), Microscopy Laboratory (RRID: SCR_017934) and Genome Technology Center (RRID: SCR_017929) for their support regarding preparation of tissue sections, microscopy, and sequencing.

## Funding

This study was supported by the National Institutes of Health/National Institute on Deafness and Other Communication Disorders (R21 DC020993).

## Data availability

All other data related to this study are available from the corresponding author upon reasonable request.

## Ethics declarations

### Competing interests

The authors declare no competing interests related to this work.

### Contributions

**Ryosuke Nakamura**: Conceptualization, Supervision, Data curation, Formal analysis, Funding acquisition, Investigation, Methodology, Project administration, Validation, Visualization, Writing – original draft. **Renjie Bing**: Data curation, Investigation, Project administration, Validation.

**Hannah Weber**: Methodology, Writing – review and editing. **Gary Gartling**: Investigation, Project administration. **Masayoshi Yoshimatsu**: Investigation, Project administration. **Michael Garabedian**: Methodology, Writing – review and editing. **Ryan Branski**: Supervision, Writing – review and editing.

## Notes

Funding for this work was provided by the National Institutes of Health/National Institute on Deafness and Other Communication Disorders (R21 DC020993)

### Competing Interest Statement

The authors have declared no competing interest.

