## Supplementary figures and images for "YAP/TAZ inhibition refines TGF-β signaling to prevent laryngeal fibrosis"

### Fig. S1

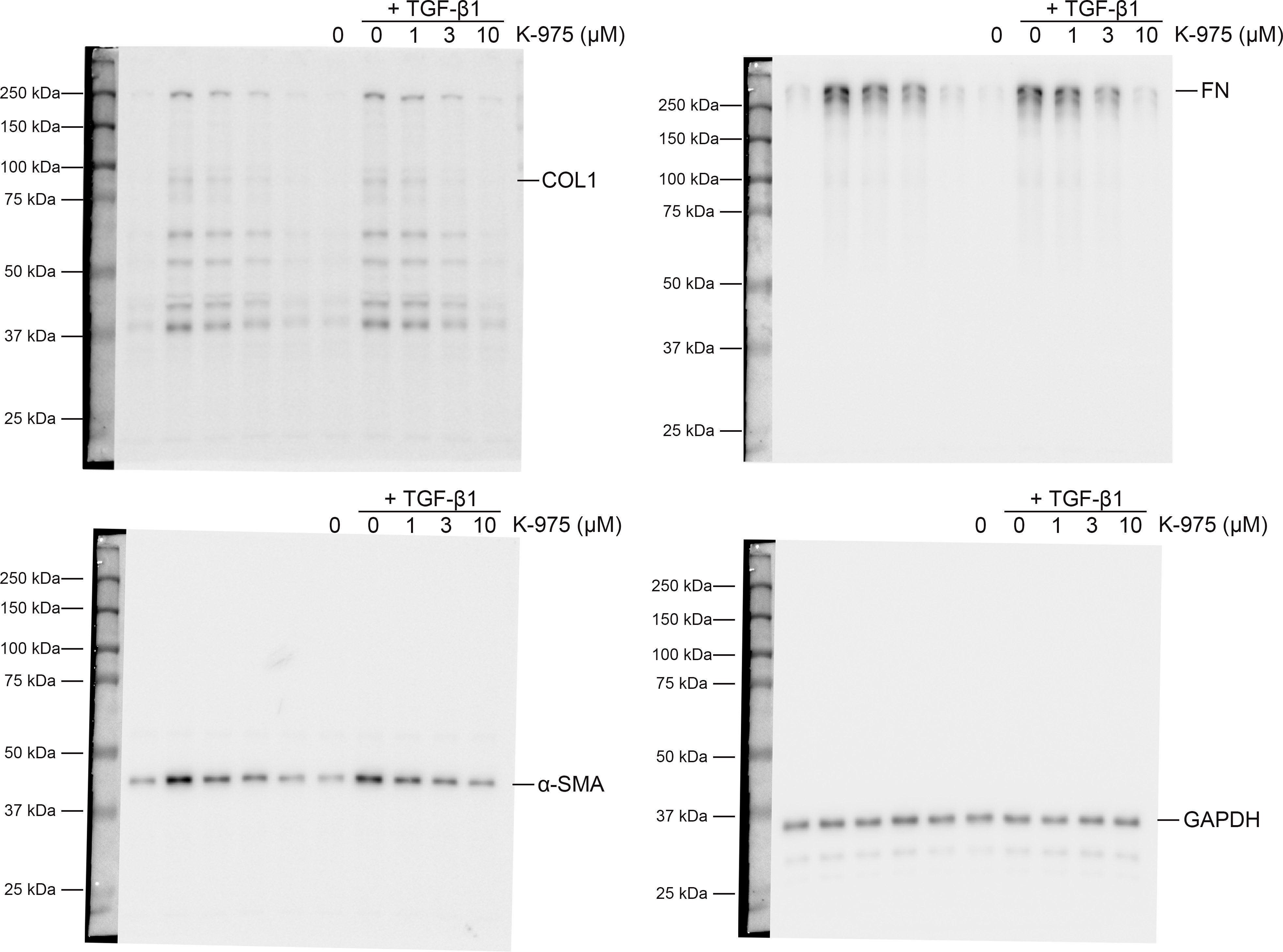

### Fig. S2

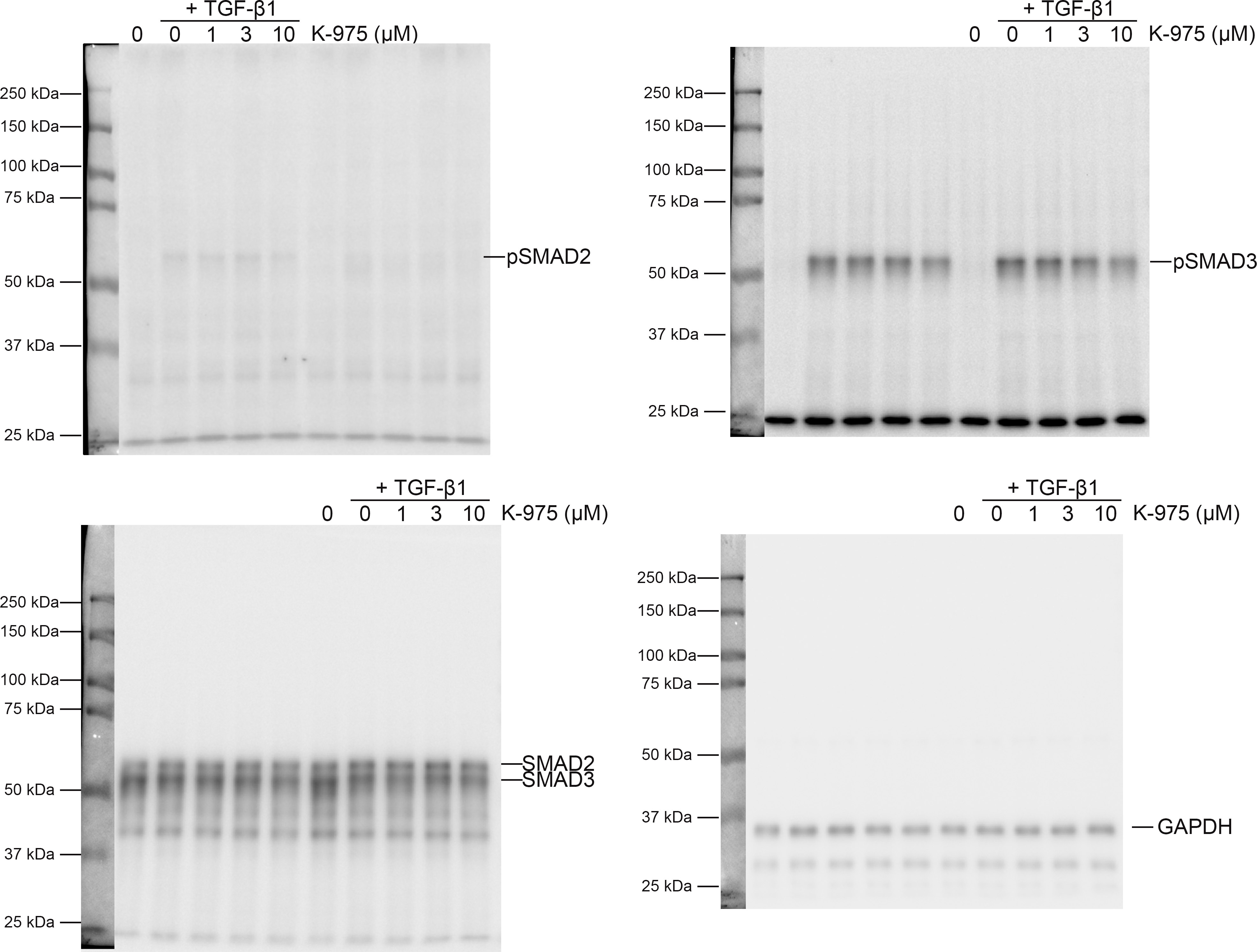
